# Coordinates and Intervals in Graph-based Reference Genomes

**DOI:** 10.1101/063206

**Authors:** Knut D. Rand, Ivar Grytten, Alexander J. Nederbragt, Geir O. Storvik, Ingrid K. Glad, Geir K. Sandve

## Abstract

**Motivation:** It has been proposed that future reference genomes should be graph structures in order to better represent the sequence diversity present in a species. However, there is currently no standard method to represent genomic intervals, such as positions of genes, on graph-based reference genomes.

**Results:** We formalize offset-based coordinate systems on graph-based reference genomes and introduce a method for representing intervals on these reference structures. We show the advantage of our method by representing genes on a graph-based representation of the GRCh38 version of the human genome and its alternative loci for regions that are highly variable.

**Conclusion:** More complex reference genomes, containing alternative loci, require methods to represent genomic data on these structures. Our proposed notation for genomic intervals makes it possible to fully utilize the alternative loci of GRCh38 and potential future graph-based reference genomes. We illustrate our notation for genomic intervals, as well as the offset-based coordinate systems, through a web tool at: https://github.com/uio-cels/gen-graph-coords.

## Introduction

A reference genome for a species makes it possible to represent genomic features from different sources in a common reference. This enables the analysis of the relationship between these features, e.g. computing the distance between a gene and a regulatory element. In order to perform such computations generically, a common coordinate system on the reference genome is needed.

Formally, a *reference genome coordinate system* is a system that uses coordinates to uniquely determine the positions of bases in the reference genome. Until recently reference genomes have been represented in linear form, meaning that there is only one path from the beginning of each chromosome (or sequence element) to the end. This enables a coordinate system where each base can be uniquely identified by the chromosome ID and the offset from the beginning of the chromosome. An example of a coordinate is chr14, offset 150000000, written more compactly as chr14:150m. A genomic interval can then be represented unambiguously by two such coordinates, the start and end position of the interval.

A problem with linear reference genomes is that they are unable to represent variation within a species, making them incapable of adequately representing features from individuals that are very different from the reference. A solution to this problem is to use *sequence graphs* as reference structures [1, 2, 3].

The newest human reference genome GRCh38 contains 261 alternative loci, regions where the human genome has been found to contain significantly different sequences in some individuals. By treating these alternative loci as side paths in a graph, GRCh38 can be seen as a sequence graph. Generally, a sequence graph can represent genomes from one or more individuals along with additional variation data. Each vertex in the graph represents a DNA-sequence of one or more base pairs, and each edge connects two consecutive sequences in the genome. Examples of the use of sequence graphs are De-Bruijn graphs in *de novo* assembly [4, 5, 6], the software project VG [7] that has built a framework for representing variation data on graphs, as well as certain applications for genotyping [8] and multiple sequence alignment [9].

Referencing positions and intervals on graph-based reference genomes poses challenges not present with linear reference genomes. First, positions such as chr1:100 will be ambiguous if there are multiple positions on chromosome 1, on different paths, having offset 100. Second, intervals represented by only a start and an end coordinate will be ambiguous, since there can be different paths taken within the interval. For instance, there is no standard method of representing genes that are partly on alternative loci in GRCh38.

In this article we address these issues by discussing suitable coordinate systems and proposing an interval representation for graph-based reference genomes, using GRCh38 and its alternative loci as an example.

## Results

### Coordinate systems on graph-based reference genomes

As mentioned, the newest human reference genome, GRCh38, can be seen as a graph. Defining a coordinate system on such graph-based reference genomes has been discussed by Marschall et al. [2]. They propose that nearby bases should have similar coordinates (*spatiality*), that coordinates should be concise and interpretable (*readability*), and that coordinates should increment along the genome (*monotonicity*). In addition, we propose that the coordinate system should be *backward compatible*, meaning that if the reference graph has been updated, coordinates from a previous version of the graph should preferably still be valid and unambiguously refer to the same bases in the new graph.

Spatiality and readability are useful in order to manually check the validity of results obtained from a computer analysis, while backward compatibility removes the need of updating all previous data when updating the reference graph. An update to the reference genome can be any number of alterations to the graph, but here we will only look at updates in which new paths are added to the graph.

In this article, we discuss a class of coordinate systems similar to the coordinate systems used on linear reference genomes, which we denote as *offset-based coordinate systems*. In offset-based coordinate systems, coordinates consist of a region identifier and an offset that is counted from the start of the region. Offset-based coordinate systems include those used on linear genomes today, with chromosome IDs as region identifiers and offsets counted from the beginning of each chromosome, e.g chr1:100. This class of coordinate system has intuitive and readable coordinates, and two coordinates representing bases close to each other in the genome will be similar. Also, computing the distance between two coordinates within the same region is as simple as taking the difference between the offsets.

Offset-based coordinate systems can be defined on graph-based reference genomes by partitioning the graph into a set of non-overlapping linear sequences, which we denote as *region paths*. There are different ways to divide the reference genome into region paths, and this is what separates different offset-based coordinate systems from each other. Here we will discuss two variants, which we refer to as *hierarchical* and *sequential partitioning*.

### The hierarchical partitioning

As mentioned, the most common way of referencing positions on GRCh38 today is using an offset-based coordinate system. This coordinate system is obtained by what we define as hierarchical partitioning: choosing one main region path through the graph, and defining alternative loci as alternative region paths.

Thus, the offsets for positions on the consensus paths are counted from the beginning of each chromosome, and offset for positions on alternative loci are counted from the beginning of each alternative locus. This can be extended to graphs with more layers of alternative loci (e.g. alternative loci of alternative loci) by choosing a main path for each layer, and defining its alternative loci as region paths.

**Figure 1:**
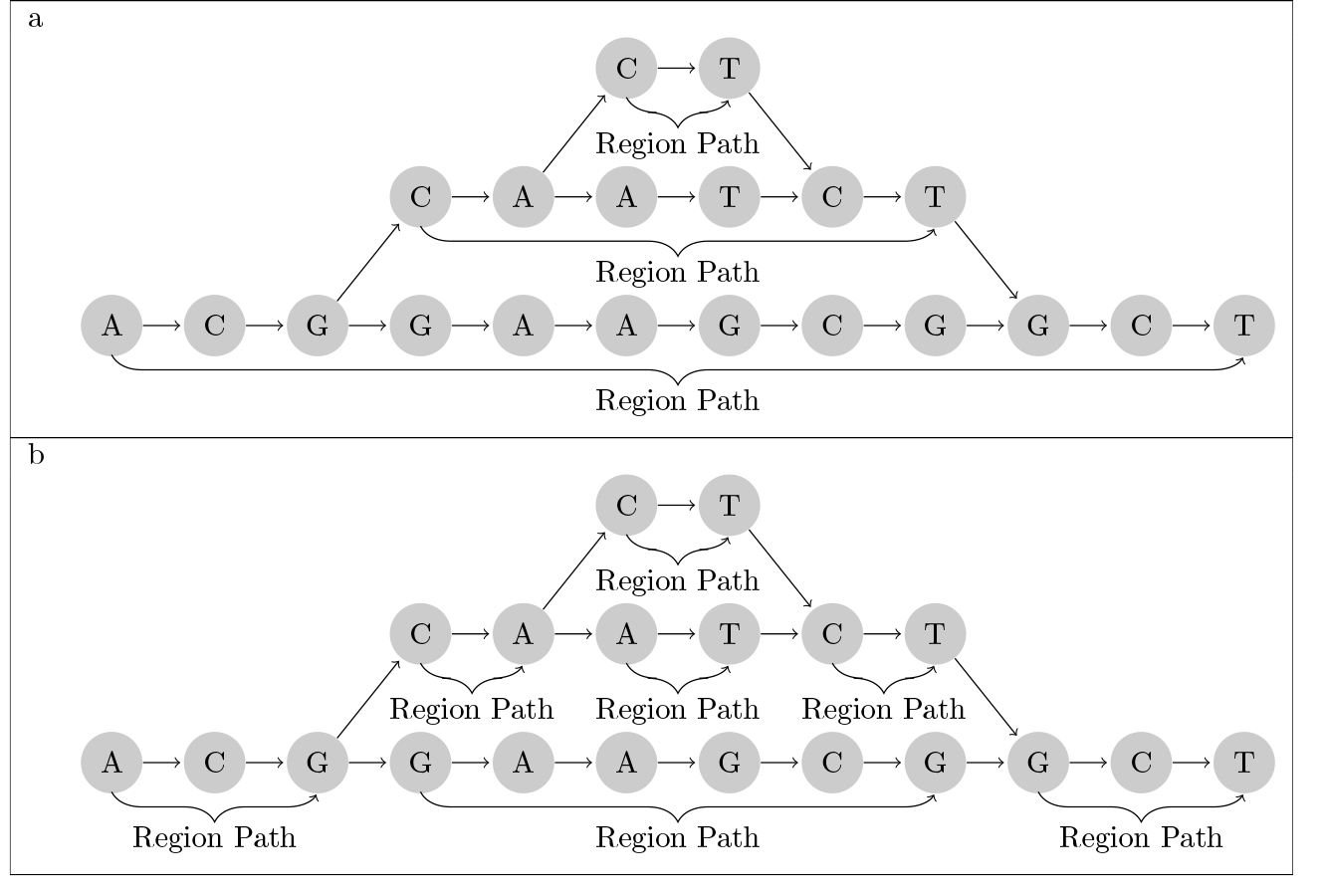
*Region path partitions*. (a) Hierarchical partitioning. The graph is partitioned into the main region path (bottom) and an alternative region path (middle) with its own alternative region path (top). (b) Sequential partitioning. The graph is partitioned into unambiguous region paths. Both the top and bottom layer are divided into three region paths.

This coordinate system is backward compatible, since adding a new path to the reference will not change the existing coordinates. However, spatiality is not fulfilled (Figure 2).

### The sequential partitioning

An alternative to the hierarchical partitioning is to also divide the main paths wherever an alternative path starts or stops (see Additional File 1 for details). Thus, adding a new alternative path to the reference graph will lead to three new region paths on the main path (before, parallel to, and after the alternative path) (Figure 1 b). This change in the partitioning breaks backward compatibility, since coordinates on the old region path are changed when updating the graph. In order to fulfill backward compatibility, one needs to keep record of what region path the three new region paths come from, which will make it possible to map coordinates on the old region path to coordinates on the new one.

**Figure 2:**
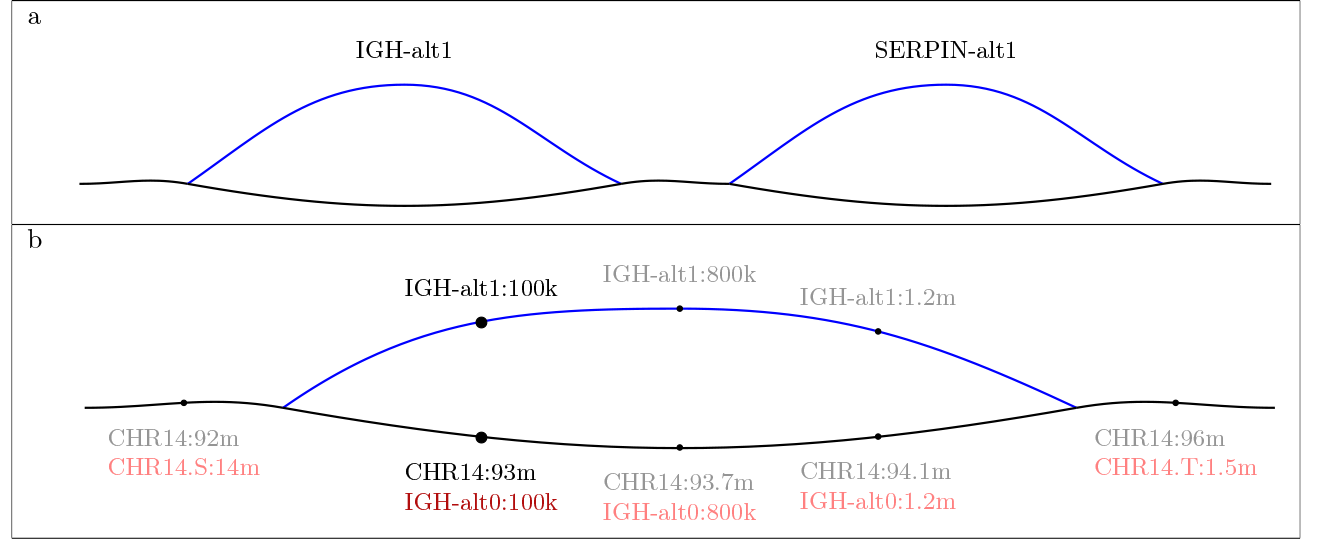
*Coordinates in GRCH38.* (a) Visualization of alternative loci in the IGH and the SERPIN region on chromosome 14 in GRCh38. The main path is shown in black and an alternative locus in blue. (b) A closer look at coordinates in the IGH region, with coordinates obtained from hierarchical partitioning (black) and sequential partitioning (red). On the alternative loci, the coordinates are the same, as this is a separate region path in both partitionings. Using the sequential partitioning, coordinates on the main path are more similar to those on the alternative loci.

## Naming Schemes

Naming the individual region paths is not straightforward, as there are multiple criteria that can be in conflict with each other: The ability to deduce from the names whether two region paths are variants of each other (vertical spatiality), whether they are close to each other along the genome (horizontal spatiality), and which comes first in the genome (monotonicity). Two naming schemes are commonly used in GRCh38: the GenBank sequence accession number (e.g. KI270758.1) [10], and a combination of the region name and the assembly-unit (e.g. MHC ALT_REF_LOCI1). The first naming scheme fulfills none of the above criteria, whereas the second only fulfills vertical spatiality.

Achieving all three criteria can be a challenge when new region paths are added. For instance, when a region path is added between two existing region paths, the name of the new region path should ideally indicate that it is closer to each of the other two region paths than they are to each other, something that can result in less readable region path names after many iterations of changes to the reference genome. In some cases, it is possible that the best solution is a naming scheme not fulfilling all criteria; sacrificing efficiency in updating the reference graph and ease of manually inspecting data for a more lightweight coordinate system.

## Interval representations on graph-based reference genomes

We define a genomic interval in a graph-based reference genome as a path between two vertices. Such intervals should be represented unambiguously. When using a linear reference structure, unambiguity can be satisfied by storing the start and end coordinates of the interval. On a graph-based reference genome, this will not give unambiguous intervals if there is more than one path between a start and an end coordinate in the graph.

To resolve this ambiguity, an interval representation must indicate which path the interval follows in the graph. The minimal solution is to indicate which region paths the interval follows wherever it is ambiguous. An interval can then be represented as a list containing the start and end coordinates in addition to information about the region paths followed by the interval.

An example of a genomic interval represented this way is (chr14:100, IGH-alt1, SERPIN-alt1, chr14:150m), which denotes the interval from offset 100 to 150m on chromosome 14 going through the region path IGH-alt1 and the region path SERPIN-alt1. This representation does not indicate that the interval follows the main path between the IGH-alt1 and the SERPIN-alt1 region paths, since the path taken here is unambiguous.

However, we argue that also including information about unambiguous paths will improve readability (see Additional File 1). In the above example, this would make it possible to deduce that the interval follows the main path in between the IGH and the SERPIN regions. The interval would then be represented as (chr14:100, IGH-alt1, chr14, SERPIN-alt1, chr14:150m). This representation is equivalent to including a region path identifier every time the identifier changes. If the coordinate system is backward compatible, intervals defined on the coordinate system will also be backward compatible, since the intervals can be uniquely determined by a set of (backward compatible) coordinates (e.g. the coordinate of the first vertex in each region path).

**Figure 3:**
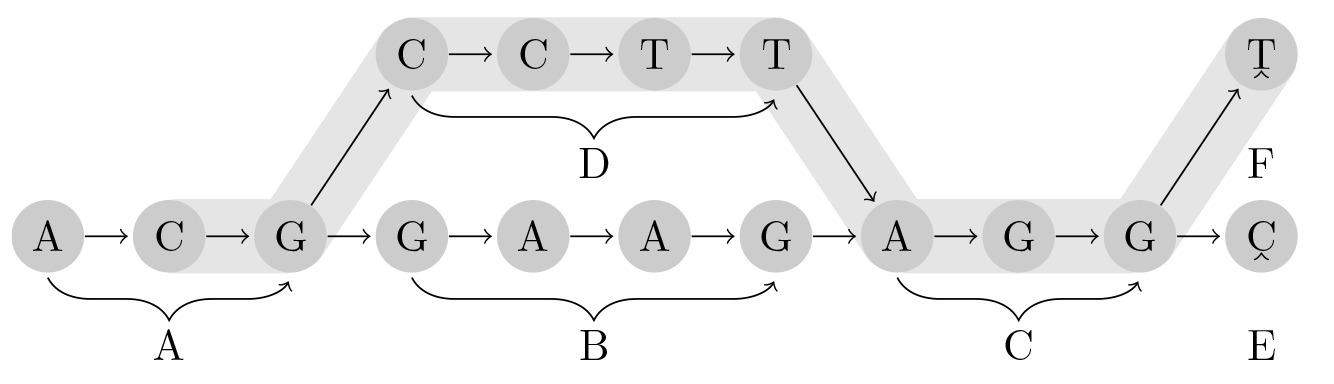
*An interval* spanning the region paths ***A, D, C*** and ***F***. Using the minimal notation, this would be represented as (***A***:1 ***D F***:0), but with the proposed full notation the ***C*** region path is also included: (***A***:1 ***D C F***:0).

## Implementation

In order to illustrate our proposed interval notation as well as the the difference between hierarchical and sequential partitioning, we have created an easy-to-use web tool that creates a graph-based reference genome based on GRCh38.

The tool uses blast [11] to align the sequence of an alternative locus in a given region with the sequence of the main path. A graph is then created by merging the sequences of the highest scoring alignments, leaving only the parts with high diversity as separate region paths in the graph.

The resulting graph can be explored through an interactive visualization, where the user can see coordinates of positions and notation of genomic intervals. Figure 4 shows an example generated by this tool, illustrating how our proposed notation for genomic intervals can be used. This tool, and its source code, can be found at https://github.com/uio-cels/gen-graph-coords.

**Figure 4:**
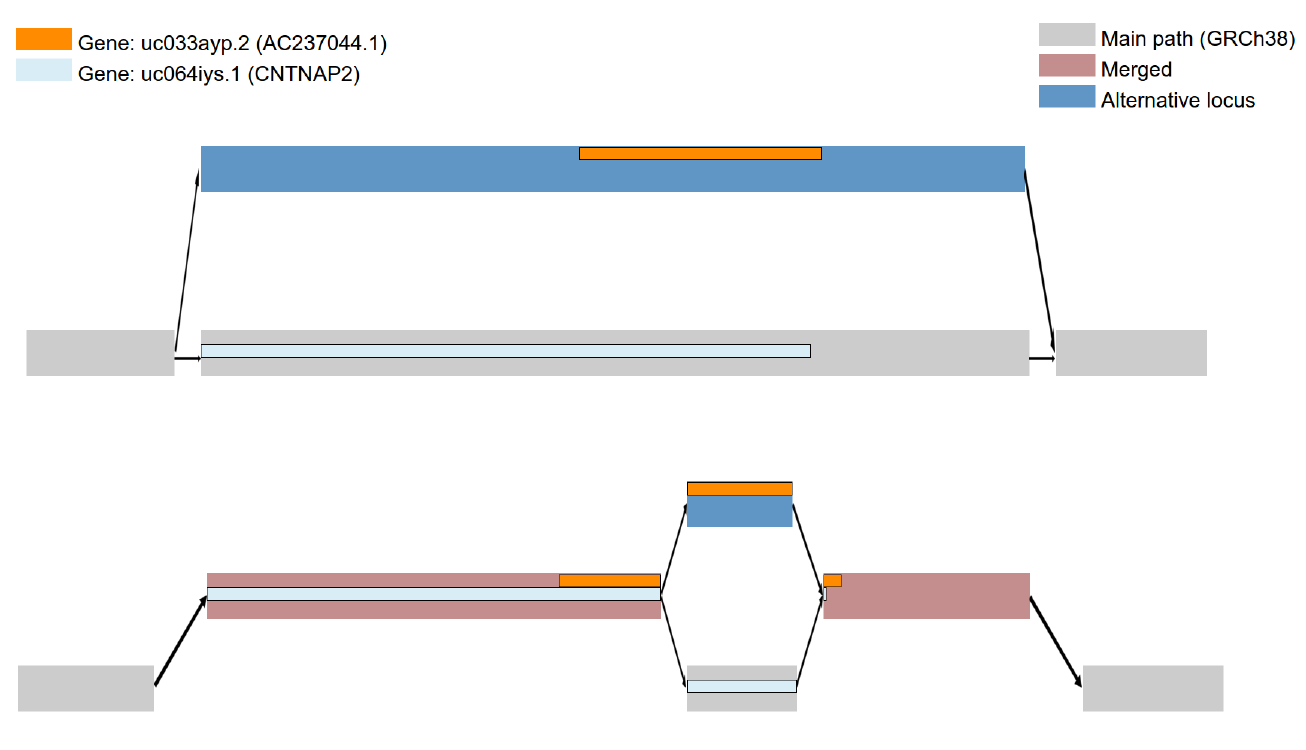
*Example from the web tool*. Visualization of two sequence-graphs, both around REGION148 on chromosome 7 from GRCh38, created by the web tool. A simple graph with the alternative locus as a side path is shown at the top. On the bottom, a graph created by merging the sequences of the highest scoring alignments is shown. Two genes (CNTNAP2 on the main path and AC237044.1 on an alternative locus) are shown. On the top sequence graph, these genes are on separate paths. After merging similar sequences, we see that AC237044.1 starts and ends on areas that have been merged with the main path. This makes it possible to analyse this gene with epigenomic data from the main path of GRCh38. Representing this gene on this graph is made possible using our genomic interval representation.

## Discussion and conclusion

After more than two years since the release of GRCh38, only few bioinformatic tools are using its alternative loci, viz. BWA-MEM [12], iBWA [13], GSNAP [14], and SRPRISM [15]. In order to utilize the potential of this additional data in GRCh38, one needs a common framework for referencing positions and intervals in the reference genome. An offset-based coordinate system makes it possible to reference positions, but there is currently no standard approach to reference intervals in graph-based coordinate systems.

We propose a simple way to unambiguously represent genomic intervals by including information about all region paths covered by the interval, as well as the start and end coordinates, using an offset-based coordinate system. Being able to represent genomic intervals on graph-based reference genomes makes it possible, for instance, to analyse a gene on an alternative locus in GRCh38 with methylation data from the main path.

Working with graph-based genomes will inevitably lead to complications not present with linear reference genomes. While the coordinate system for linear reference genomes is simple and achieves our discussed criteria (spatiality, readability, monotonicity and backward compatibility), a graph-based coordinate system will be much more complex and only partially meet these criteria. Thus it is necessary to weigh the different criteria, as well as the overall goal of simplicity, against each other in order to find the most suitable system.

With a system for representing genomic intervals on graph-based reference genomes and a common coordinate system, researchers can begin to utilize more of the full potential of GRCh38 and future graph-based reference genomes.

## Availability of data and material

Data on alternative loci and genes in GRCh38 were collected from the public USCS MySQL database at genome-mysql.cse.ucsc.edu. The sequences used for alignment were collected using the togows.org API.

Our software project “Gen-Graph-Coords” can be found at https://github.com/uio-cels/gen-graph-coords, version 1.0. The project is platform independent and programmed in *Python, Javascript* and *HTML*. Licensed under the GNU PL license.

## Author’s contributions

IG and KDR drafted the manuscript, and developed the software. All authors took part in manuscript writing and read and approved the final version of the manuscript.

## Additional Files

### Additional file 1 — Definitions

Formal definitions of terms used in the article (pdf).

